# Disrupting action control with transcranial ultrasound neuromodulation: a step forward for Tourette syndrome

**DOI:** 10.1101/2025.10.29.685348

**Authors:** Cyril Atkinson-Clement, Mairi Houlgreave, Aikaterini Gialopsou, Caitlin Mairi Smith, Isabel Farr, Aneta Dvorakova, James Kennaway, Stephen R. Jackson

**Author notes:** **Corresponding author:** Dr. Cyril Atkinson-Clement.

## Abstract

**Background:** While Tourette syndrome (TS) is characterised by the presence of tics, premonitory urges are involved in both triggering them and as a lever to control them. The right insular cortex is known to play a key role in these processes, yet its precise functional contribution remains unclear. Transcranial ultrasound stimulation (TUS) is currently one of the only non-invasive neuromodulation techniques capable of safely and precisely modulating the activity of deep brain structures such as the insula.

**Objective:** This study used TUS to causally investigate the respective roles of the anterior and posterior insula in urge and action control during a blink suppression task.

**Methods:** Twenty healthy adults underwent three TUS sessions targeting the right anterior insula, posterior insula, and posterior ventricle (control site) in a within-subject, crossover design. Behavioural outcomes included blink counts and urge ratings. Additionally, voxel-based correlations were conducted to further estimate the role of each reached area with acoustic stimulation, allowing for a better mechanical understanding.

**Results:** TUS of the anterior insula induced a clear and acute increase in blink frequency without altering urge, reinforced by voxel-based analyses, confirming its role in voluntary action control. Posterior insula stimulation induced delayed increases in both blinks and urge, which are essentially explained by an involuntary stimulation of the right inferior frontal gyrus. Ventricular stimulation unexpectedly reduced blink frequency during the stimulation only, correlated with acoustic energy delivered within the posterior internal capsule. No significant side effects or temperature rises were observed.

**Conclusions:** These findings provide the first causal evidence that TUS can modulate the two main features of TS: premonitory urge and action control. Distinct effects across targets highlight the anterior insula’s critical role in action control and suggest the internal capsule as a potential therapeutic target for TS.

## Introduction

Tourette syndrome (TS) is a neurodevelopmental condition characterised by the presence of motor and vocal tics [1]. Tics are often preceded by uncomfortable sensations known as premonitory urges, which are transiently relieved by performing the tic [2]. These urges play a central and dual role in TS, being both the problem and perhaps the solution. If they frequently act as the subjective trigger for tic expression, they can also serve as a key focus for behavioural interventions. Urges make TS the only movement disorder for which the majority of patients can have voluntary action control for varying durations [2,3]. Comprehensive behavioural intervention for tics, for example, aims to enhance patients’ awareness of premonitory urges and to disrupt the tic expression cycle through voluntary inhibition. However, while it has demonstrated good efficacy, a substantial proportion of individuals with TS show insufficient improvement [4], highlighting the need for additional clinical approaches, including pharmacological treatment and, in severe cases, deep brain stimulation.

From a neural perspective, premonitory urges involve a network encompassing the insular cortex, anterior and mid-cingulate cortices, and the parietal operculum [5,6]. Many investigations of premonitory urges have been conducted in healthy participants using paradigms such as blink or yawn suppression, which elicit sensations comparable to the urge-to-tic experienced by individuals with TS [6–8]. A recent meta-analysis indicated that the right insular cortex is consistently engaged across various urge-related paradigms, including blink suppression, and is also part of the broader TS network [9]. Moreover, functional differentiation within the insula has been observed: the urge-to-blink appears to recruit both anterior and posterior insular regions, whereas blink suppression primarily involves the anterior insula [10,11], pointing to distinct functional roles within the insular cortex that remain to be clarified.

Recent advances in non-invasive brain stimulation techniques, and especially in transcranial ultrasound stimulation (TUS) have opened new opportunities to investigate causal relationships between specific brain regions and behavioural processes, an achievement that was previously impossible without invasive methods. TUS enables the safe, painless, and transient modulation of both cortical and deep brain regions with high spatial precision [12–17]. In practice, TUS consists of delivering multiple focused acoustic ultrasound waves to a specific brain location, where the combined acoustic energy can transiently alter neural activity [14,18]. Although its precise mechanisms remain under debate, TUS effects have been linked to mechanical modulation of ion channels [19], micro-cavitation depolarisation [20] and altered neuron-glia coupling [21]. The number of studies using TUS has rapidly increased, demonstrating its potential as both a research and therapeutic tool across a range of clinical populations [22].

The present study aimed to examine the specific contributions of the anterior and posterior insular cortices on the urge-to-blink and action control. To this end, healthy volunteers performed a blink suppression task while TUS was applied to either the anterior or posterior parts of the right insula, with a control condition targeting the posterior ventricle. The behavioural paradigm was designed to capture both immediate and delayed (up to 30 minutes) effects of TUS. Findings from this study are expected to inform future work exploring the causal role of insular subregions in TS.

## Methods

### Participants

We recruited 20 healthy and right-handed volunteers with no history of neurological or psychiatric disorders (see Table.1). Exclusion criteria included: a family history of seizures, predisposition to syncope, excessive hair that could interfere with ultrasound transducer coupling, current or planned pregnancy, implanted metallic devices, dermatological conditions affecting the scalp, claustrophobia or anxiety related to MRI, and tattoos located near the head.

Participants were recruited through university mailing lists, advertisements, and personal contacts, and they received £30 compensation upon completing the study.

The study was approved by the Ethics Committee of the University of Nottingham, Faculty of Psychology (reference: F1298R), and conducted in accordance with the revised Declaration of Helsinki [23]. All participants provided written informed consent before their participation.

### Study design

Following recruitment, a T1-weighted structural MRI scan was acquired for each participant using a 3 Tesla scanner (Philips Ingenia; matrix size=256*256, field of view=176*256*256mm, voxel size=1mm isotropic, 176 slices, TR/TE=8.1/3.7ms, flip angle=8°) at the Sir Peter Mansfield Imaging Centre, University of Nottingham, unless participants provided a recent scan. This sequence was used to plan the TUS sessions, which involved: i) identifying the target coordinates, ii) identifying the optimal TUS transducer position, iii) estimating a T1-based pseudo-computed tomography (PCT), iv) segmenting the tissues to identify the skull, and v) running acoustic simulations to estimate the delivered acoustic energy and temperature increase (see supplementary materials).

Each participant completed three experimental visits, each lasting approximately one hour and scheduled at least one week apart. To minimise intra-subject variability, sessions were arranged at approximately the same time of day (within 90 minutes) and preferably on the same weekday. Across the three visits, participants received TUS targeting one of the three sites: the right anterior insula (TUS-AntIns), the right posterior insula (TUS-PostIns), or the right posterior lateral ventricle (TUS-Vent). The order of target stimulation was counterbalanced across participants.

At the start of each visit, participants completed a baseline questionnaire to assess potential side effects, rating any physical sensations on a scale from 0 (“absent”) to 10 (“maximal”). Neuronavigation was then calibrated (Brainsight; Rogue Research, Montreal, Canada), and the TUS transducer was positioned at the predetermined entry point, aligned with the target. Electrooculography (EOG) electrodes were placed above and below the left eye, with the ground electrode placed on the back of the left wrist. EOG signals were recorded at a sampling rate of 30kHz using Signal acquisition software (version 8.03, Cambridge Electronic Design, Cambridge, UK). Participants wore bone-conduction headphones that produced a sounds that imitated TUS [24] (available here: https://github.com/benjamin-kop/auditory-masking-stimuli/tree/1).

After the setup, participants performed the 45-minute behavioural task. Once completed, all devices were removed, and participants washed their hair. After this, they filled out the side-effect questionnaire again. To evaluate potential delayed effects, participants were asked to complete the side-effect questionnaire once more on the following day.

### TUS planning

Brain target coordinates were identified using either Montreal Neurological Institute (MNI) coordinates for the insular targets or each participant’s native T1-weighted scan for the ventricular target. Insular targets were defined using the AAL atlas [25], where the right insula was manually subdivided into anterior and posterior portions of equal voxel count in the MNI space (see supplementary materials). The centroids of these subdivisions were used as target coordinates (anterior insula: x=38, y=16, z=-3; posterior insula: x=40, y=-8, z=6). MNI coordinates were then individually transformed to the native space using the registration tools provided by Advanced Normalization Tools (ANTs) [26].

Transducer positioning was performed using a dedicated, validated and open-source code designed to minimise the angle between the transducer and the head [27]. It starts by selecting all scalp coordinates within a distance that the transducer can reach from the target (neither too far nor too close). Then, it estimates the volume between the head and the transducer exit plane and identifies the coordinate with the lower value as the optimal one. Outputs from this process are used to set up the neuronavigation (see Figure. 1), determine the target depth, and perform acoustic simulations.

**Figure. 1.**
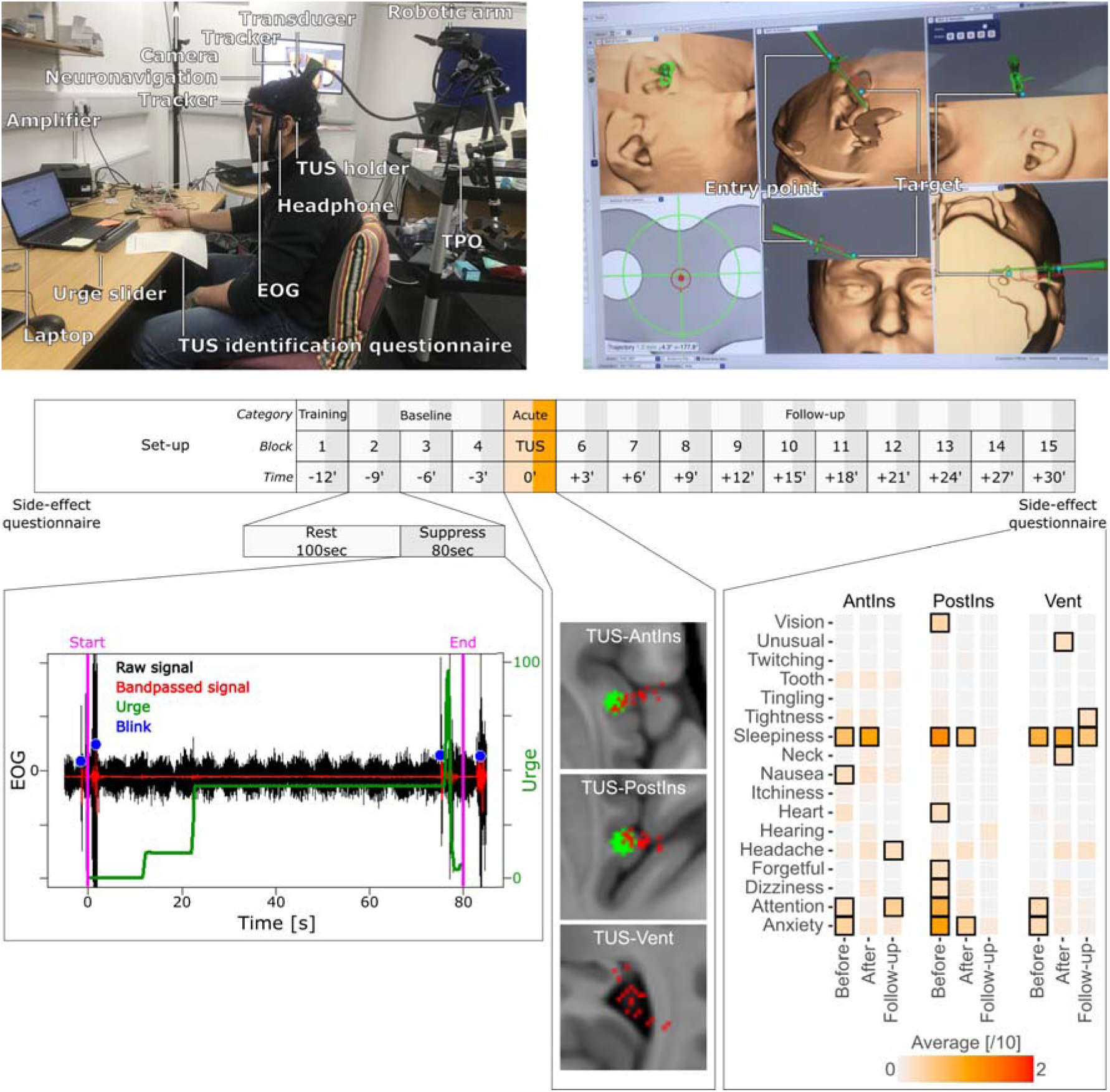
Keys steps of the experimental protocol. Top panels illustrate the experimental setup and the various devices used (left) and demonstrates how the target was reached by positioning the transducer at the previously estimated entry point (right). Panel below shows the task composition, as well as the EOG preprocessing which enables the automatic detection of blinks along with its relation to the dynamic urge score, the locations of peak pressure for the 20 participants across the three targets, and the side effects measured before the procedure, immediately after, and 24 hours post-TUS. The highlighted squares indicate side effects that were reported significantly (with an averaged value that was significantly higher than 0).

### Experimental setup

On each experimental day, participants were instructed to remain seated in a chair while the experimental devices were connected (see Figure. 1). First, EOG electrodes were placed above and below the left eye, with the ground electrode positioned on the back of the left hand. Next, bone-conduction headphones were positioned, and the neuronavigation tracker was attached to the participant’s forehead.

Degassed ultrasound gel was then applied both to the participant’s scalp at the identified transducer position and to the transducer itself. The transducer was carefully positioned and oriented according to the target coordinates and secured to the participant’s head to maintain immobility throughout the session. If necessary, the transducer position was adjusted during the session to ensure accurate targeting. This procedure ensured that the transducer remained in a fixed position throughout the entire 45-minute experiment, it allowed the participant to make slight movements during that time, and kept them unaware of when the device was active or inactive.

### Behavioural task

The behavioural task was implemented in Matlab Psychtoolbox (http://psychtoolbox.org) and lasted 45 minutes, consisting of 15 blocks of 3 minutes each. Each block was divided into two parts (see Figure. 1).

During the first part (100 seconds), participants were free to move and blink while remaining seated. A countdown was displayed on the laptop screen indicating the remaining time before the second part. In the second part (80 seconds), participants were instructed to refrain from blinking while continuously rating their urge to blink using a slider. During this period, the word “suppress” was displayed on the screen, while a noise was delivered through bone-conduction headphones, and blink activity was recorded via the EOG system.

After each block, participants reported whether they believed TUS had been applied using a scale from −5 (certain stimulation was off) to +5 (certain stimulation was on), with 0 indicating uncertainty. TUS was applied exclusively during block 5, allowing for an initial familiarisation block (block 1; discarded in statistical analyses), three baseline blocks (blocks 2–4), one block measuring the acute TUS effect (block 5), and ten follow-up blocks (blocks 6–15) spanning over 30 minutes.

All experimental elements (including EOG and urge recording, white noise, and TUS delivery) were controlled through the Matlab code to ensure precise synchronisation.

### TUS application

TUS was delivered during block 5 using a four-element CTX-500-4CH transducer coupled with the NeuroFUS PRO TPO-203 system (Sonic Concepts, Brainbox Ltd., Cardiff, United Kingdom). The stimulation protocol was identical for all participants and targets, with the following parameters: central frequency=500kHz; spatial peak pulse average intensity (Isppa) in water=54.51W/cm²; pulse duration=20ms; pulse repetition interval=200ms; pulse repetition frequency=5Hz; duty cycle=10%; total stimulation duration=80s.

Degassed ultrasound transmission gel was applied to both the participant’s scalp and the transducer face to ensure effective coupling.

### TUS acoustic simulation

Acoustic simulations were performed using k-Plan v1.2 on each participant’s T1-weighted structural scan and the corresponding PCT derived from it [28]. The resulting maps of estimated acoustic pressure and temperature rise (in °C) were transformed from native space to template space using ANTs. Peak pressure, peak temperature rise, and the transcranial mechanical index (MItc) were calculated within brain tissue and are summarised in the Table 1. Figure. 1 illustrates the peak pressure locations for each participant relative to the brain targets.

**Table. 1.** Demographic and acoustic properties for the three TUS targets.

|  | Gender | Age (y) | TUS-AntIns |  |  |  | TUS-PostIns |  |  |  | TUS-Vent |  |  |  |
| --- | --- | --- | --- | --- | --- | --- | --- | --- | --- | --- | --- | --- | --- | --- |
|  |  |  | Coupling volume [ml] | Peak pressure [kPa] | MItc | +°C | Coupling volume [ml] | Peak pressure [kPa] | MItc | +°C | Coupling volume [ml] | Peak pressure [kPa] | MItc | +°C |
| ID001 | M | 27.7 | 11.06 | 546.1 | 0.785 | 0.31 | 6.12 | 619.5 | 0.879 | 0.50 | 6.83 | 564.4 | 0.806 | 0.51 |
| ID002 | F | 41.3 | 11.03 | 385.9 | 0.556 | 1.06 | 8.21 | 500.2 | 0.718 | 0.64 | 7.89 | 604.0 | 0.873 | 0.70 |
| ID003 | F | 30.8 | 13.72 | 417.1 | 0.613 | 0.29 | 7.97 | 451.6 | 0.651 | 0.39 | 10.49 | 645.3 | 0.932 | 0.37 |
| ID005 | F | 21.8 | 15.99 | 314.9 | 0.593 | 0.65 | 11.48 | 434.8 | 0.623 | 0.82 | 5.18 | 663.8 | 0.956 | 0.37 |
| ID006 | F | 20.2 | 15.86 | 525.5 | 0.762 | 0.49 | 6.43 | 571.4 | 0.817 | 1.05 | 3.78 | 592.5 | 0.856 | 1.12 |
| ID007 | F | 23.1 | 13.38 | 301.8 | 0.815 | 2.15 | 7.67 | 548.7 | 0.799 | 1.81 | 8.76 | 617.0 | 0.894 | 0.46 |
| ID008 | F | 22.3 | 9.06 | 400.1 | 0.572 | 0.29 | 4.04 | 500.5 | 0.723 | 0.44 | 3.69 | 520.3 | 0.743 | 0.75 |
| ID010 | M | 24.6 | 6.81 | 386.4 | 0.555 | 0.39 | 3.65 | 499.5 | 0.725 | 0.42 | 4.34 | 579.7 | 0.841 | 0.92 |
| ID011 | M | 25.2 | 6.20 | 370.1 | 0.537 | 0.60 | 8.02 | 504.6 | 0.735 | 0.76 | 5.47 | 660.4 | 0.955 | 1.00 |
| ID012 | M | 24.7 | 6.69 | 511.8 | 0.748 | 0.25 | 7.11 | 521.3 | 0.756 | 0.22 | 2.03 | 375.5 | 0.541 | 0.95 |
| ID013 | F | 19.2 | 7.86 | 278.8 | 0.400 | 0.53 | 3.99 | 304.2 | 0.438 | 0.19 | 6.98 | 458.7 | 0.663 | 0.18 |
| ID014 | M | 51.1 | 7.54 | 282.7 | 0.416 | 0.44 | 4.69 | 384.9 | 0.556 | 0.46 | 2.42 | 439.6 | 0.631 | 0.68 |
| ID015 | M | 29.7 | 5.27 | 516.8 | 0.734 | 0.19 | 6.59 | 454.1 | 0.650 | 0.34 | 5.41 | 562.9 | 0.814 | 0.52 |
| ID016 | M | 28.4 | 10.98 | 570.1 | 0.857 | 0.51 | 12.20 | 496.7 | 0.729 | 1.38 | 6.85 | 575.6 | 0.823 | 1.48 |
| ID019 | F | 21.9 | 16.21 | 431.4 | 0.614 | 0.42 | 11.46 | 559.1 | 0.802 | 0.28 | 6.19 | 617.1 | 0.883 | 0.44 |
| ID020 | F | 30.2 | 23.39 | 237.4 | 0.487 | 0.99 | 13.55 | 410.1 | 0.597 | 0.47 | 4.96 | 470.7 | 0.686 | 0.34 |
| ID021 | F | 26.2 | 12.44 | 300.9 | 0.535 | 0.78 | 9.39 | 545.4 | 0.778 | 0.41 | 4.92 | 513.6 | 0.738 | 0.35 |
| ID022 | F | 40.3 | 13.40 | 298.3 | 0.430 | 0.75 | 8.36 | 510.6 | 0.743 | 0.49 | 4.94 | 661.0 | 0.943 | 0.52 |
| ID023 | F | 27.0 | 15.63 | 376.9 | 0.563 | 0.31 | 9.81 | 313.2 | 0.449 | 0.46 | 4.15 | 385.0 | 0.557 | 0.34 |
| ID024 | F | 29.3 | 18.73 | 274.1 | 0.488 | 0.14 | 22.89 | 456.9 | 0.657 | 0.48 | 10.50 | 428.4 | 0.618 | 0.44 |
| Mean | 7M 13F | 28.2 | 12.06 | 386.3 | 0.603 | 0.58 | 8.68 | 479.4 | 0.691 | 0.60 | 5.79 | 546.8 | 0.788 | 0.62 |
| SD |  | 7.9 | 4.74 | 102.4 | 0.137 | 0.45 | 4.36 | 80.9 | 0.116 | 0.40 | 2.33 | 92.7 | 0.133 | 0.33 |
| Min |  | 19.2 | 5.27 | 237.4 | 0.400 | 0.14 | 3.65 | 304.2 | 0.438 | 0.19 | 2.03 | 375.5 | 0.541 | 0.18 |
| Max |  | 51.1 | 23.39 | 570.1 | 0.857 | 2.15 | 22.89 | 619.5 | 0.879 | 1.81 | 10.50 | 663.8 | 0.956 | 1.48 |

Coupling volume is provided by the TUS transducer positioning code [27]. Peak pressure, MItc and temperature increase are provided by k-Plan.
AntIns: Anterior insula; F: Female; M: Male; MItc: transcranial mechanical index; PostIns: Posterior insula; SD: Standard deviation; TUS:
Transcranial ultrasound stimulation; Vent: Ventricle.

### Statistical analysis

All statistical analyses were conducted in R (version 4.4.2). For the behavioural task, the first block was treated as a familiarisation period and excluded from analyses.

Blink counts per block were derived from the EOG signal. Preprocessing included low-pass filtering to remove frequencies above 75Hz, followed by z-scoring. A blink was defined as an event in which the absolute z-score exceeded a value of 4, with a minimum interval of 2 seconds between successive blinks (see Figure. 1).

Urge ratings were obtained by normalising slider values to a 0–100 scale, where 0 corresponded to the leftmost slider position (“no urge”) and 100 to the rightmost position (“maximal urge”). For each suppression period (80 seconds), the average urge score was computed.

For both blink counts and urge ratings, dependent measures were defined as the difference between block-specific values and the average of the three pre-TUS baseline blocks (blocks 2–4). The certainty of stimulation detection (−5 for a certainty that TUS was off, +5 for a certainty that TUS was on) was also included as a dependent measure.

Statistical analyses employed linear repeated measures mixed models with block number (14 levels), stimulation target (3 levels), and their interaction as fixed factors, while the participants’ ID were included as a repeated factor (20 levels). For models of blink count and urge score, the TUS detection rating was included as a covariate to take into account any possible placebo effect. When significant main effects were identified (p≤0.05), post-hoc pairwise comparisons were performed with false discovery rate (FDR) correction, including comparisons against baseline (difference from 0).

For significant behavioural effects, exploratory correlations were performed with MRI-based acoustic simulations. These analyses examined voxel-based associations between behavioural outcomes (change in blink count or urge score relative to baseline) and the estimated acoustic pressure. Analyses were restricted to regions of interest (ROIs) in which at least one participant received ≥100kPa during TUS. To account for variability in ROI size across targets (TUS-AntIns=13,958 voxels; TUS-PostIns=22,667 voxels; TUS-Vent=69,705 voxels), voxel-based significance was set at p≤0.001 with a minimum cluster size of 10 contiguous voxels (voxel size of 1mm^3^).

## Results

### Targets comparison

Repeated measures mixed models revealed significant differences in TUS delivery across the three targets (see Table.1). Coupling quality differed between targets (F_(2,38)_=24.005, p<0.001, Cohen’s f=1.055), with TUS-Vent showing the best coupling (5.79±2.33ml), followed by TUS-PostIns (8.68±4.36ml) and TUS-AntIns (12.06±4.74ml). Peak delivered acoustic energy also varied by target (F_(2,38)_=23.109, p<0.001, Cohen’s f=1.009), with TUS-Vent exhibiting the highest peak pressure (546.77±92.75kPa), followed by TUS-PostIns (476.36±80.95kPa) and TUS-AntIns (386.35±102.42kPa). Similarly, the MItc differed across targets (F_(2,38)_=17.172, p<0.001, Cohen’s f=0.88), being highest for TUS-Vent (0.79±0.13), intermediate for TUS-PostIns (0.69±0.11), and lowest for TUS-AntIns (0.60±0.14). In contrast, there was no significant difference in temperature rise between targets (F_(2,38)_=0.097, p=0.908, Cohen’s f=0.065). The Figure. 2 illustrates the averaged acoustic pressure for each target, as well as the areas for which at least 100kPa were delivered.

**Figure. 2.**
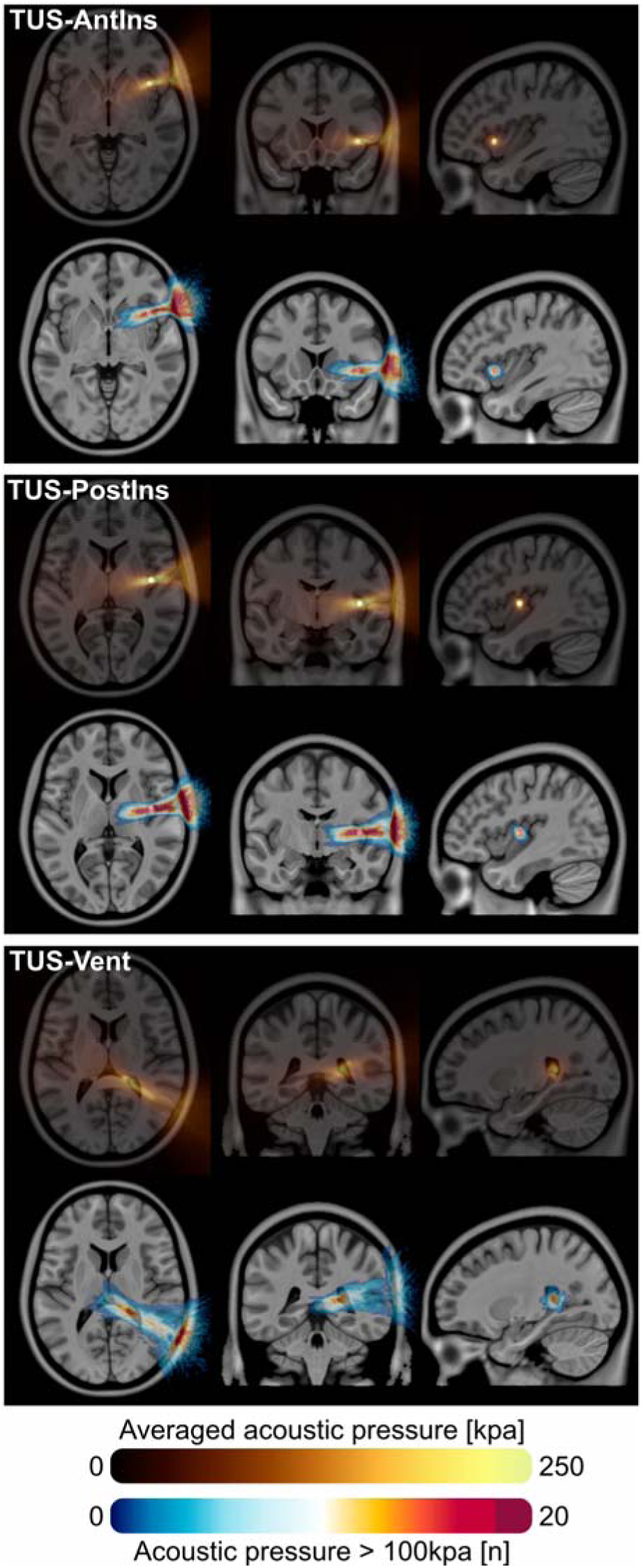
Delivered acoustic energy. For each target, the figure highlights the averaged acoustic intensity delivered (brown to light colour scale) and the number of participants who received at least 100kPa (blue to red colour scale). The targets are presented as a white sphere of radius 3mm.

Coupling quality was negatively correlated with both peak pressure (t_(58)_=-3.304, p=0.001, r=-0.398, Cohen’s f=0.434) and MItc (t_(58)_=–2.212, p=0.031, r=-0.279, Cohen’s f=0.291), but showed no relationship with temperature rise (t_(58)_=0.352, p=0.726, r=0.046, Cohen’s f=0.046).

### TUS side effects

Exploratory analyses of secondary effects (see Figure. 1) revealed no main effect of target (F_(2,2907)_=1.675, p=0.187, Cohen’s f=0.034) and no interaction between target and side-effect category (F_(32,2907)_=0.999, p=0.469, Cohen’s f=0.105). Conversely, a significant effect of target over time (delay following TUS) was observed (F_(4,2907)_=6.152, p<0.001, Cohen’s f=0.092). However, post-hoc analyses indicated that average side-effect scores were higher before TUS than after, suggesting no adverse effects induced by stimulation. No significant three-way interaction between target, delay, and side-effect category was found (F_(64,2907)_=0.747, p=0.933, Cohen’s f=0.129), further confirming the absence of TUS-related side effects. Significant effects were found for the nature of the side effects themselves (F_(16,2907)_=15.299, p<0.001, Cohen’s f=0.291) and in interaction with delay (F_(32,2907)_=2.654, p<0.001, Cohen’s f=0.717). Post-hoc analyses indicated that anxiety and attention complaints were higher before TUS than immediately after (p<0.001), with no differences observed on the day following the session (p>0.19).

### Behavioural results

Despite the use of bone-conduction headphones with white noise to mask TUS sounds, participants were often able to detect stimulation (see Figure. 3). Detection was similar across the three targets (block main effect: F_(13,798)_=3.511, p<0.001, Cohen’s f=0.248; target main effect: F_(2,798)_=0.165, p=0.848, Cohen’s f=0.021; block * target interaction: F_(26,798)_=0.890, p=0.624, Cohen’s f=0.176). FDR-corrected post-hoc comparisons indicated that the TUS identification score was significantly greater than 0 only during stimulation (block 5): TUS-AntIns (marginal mean=1.6, p=0.008), TUS-PostIns (marginal mean=1.4, p=0.013), and TUS-Vent (marginal mean=0.95, p=0.073), with no significant differences between targets (p=0.788). Based on these results, subsequent analyses included the TUS identification score as a nuisance covariate.

**Figure. 3.**
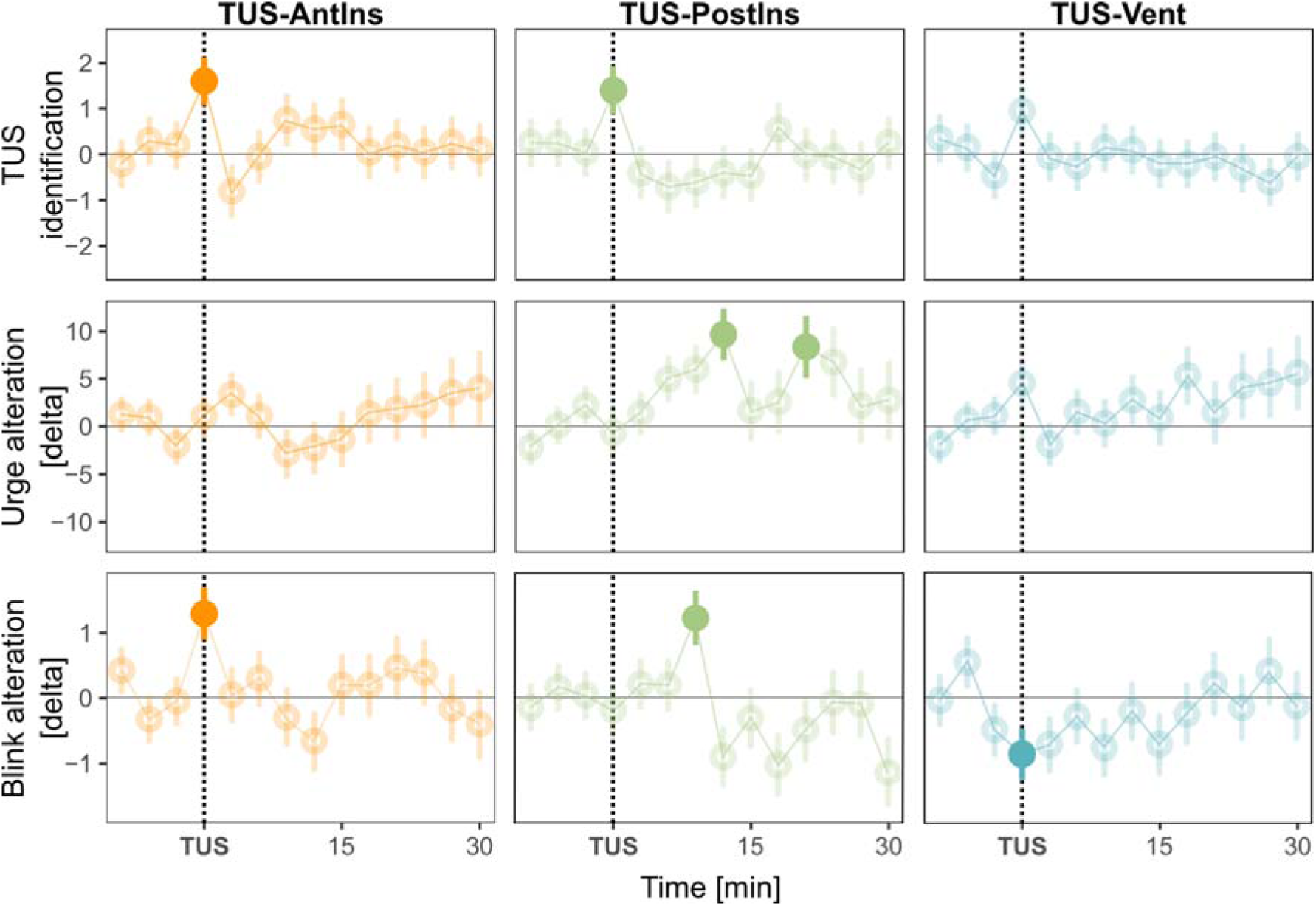
Behavioural effects of TUS. For each target, the figure highlights the averaged score of TUS identification, the averaged alteration of urge score and the number of blinks (both centred to the average of the baseline blocks, 2 to 4). Opaque points correspond to FDR-corrected significant effects.

For averaged urge ratings, no significant main effects were observed for block (F_(13,784)_=1.018, p=0.431, Cohen’s f=0.135) or target (F_(2,784)_=0.372, p=0.691, Cohen’s f=0.029; see Figure. 3), but the block * target interaction was significant (F_(26,784)_=1.919, p=0.004, Cohen’s f=0.249). This interaction was specific to TUS-PostIns, which showed increased average urge scores during block 9 (12 minutes following TUS; marginal mean=9.669, p=0.002) and 12 (21 minutes following TUS; marginal mean=8.351, p=0.038), with a trend during block 8 (9 minutes after TUS; marginal mean=6.012, p=0.062).

Similarly, for blink counts per block, no significant main effects were found for block (F_(13,784)_=1.270, p=0.226, Cohen’s f=0.151) or target (F_(2,784)_=2.343, p=0.105, Cohen’s f=0.078; see Figure. 3), whereas a significant block * target interaction emerged (F_(26,784)_=2.258, p<0.001, Cohen’s f=0.277). Post-hoc analyses revealed acute TUS effects: an increased number of blinks during TUS-AntIns (block 5; marginal mean=1.29, p=0.004) and a decreased number of blinks during TUS-Vent (block 5; marginal mean=-0.87, p=0.039). The increase during TUS-AntIns was significantly different from both TUS-PostIns (p=0.009) and TUS-Vent (p<0.001), whereas TUS-PostIns and TUS-Vent did not differ significantly (p=0.21). Nine minutes after TUS-PostIns (block 8), blink counts increased (marginal mean=1.22, p=0.01), differing significantly from TUS-AntIns (p=0.016) and TUS-Vent (p=0.002).

### Brain correlates

Voxel-based analyses revealed significant relationships between behavioural outcomes and delivered acoustic energy only for blink count changes (see Figure. 4). For TUS-AntIns, the increase in blinks observed during stimulation was positively correlated with acoustic pressure in a cluster of 21 voxels localised in the anterior insula (peak coordinates: x=31, y=18, z=0; peak r=0.779, p=0.00005). For TUS-PostIns, the increase in blinks observed nine minutes after TUS was negatively correlated with a cluster of 93 voxels in the inferior frontal gyrus (IFG; peak coordinates: x=61, y=8, z=11; peak r=–0.818, p=0.00001). For TUS-Vent, the decrease in blinks during stimulation was associated with a cluster of 40 voxels in the posterior limb of the internal capsule (PLIC; peak coordinates: x=36, y=–38, z=5; peak r=– 0.75, p=0.00014).

**Figure. 4.**
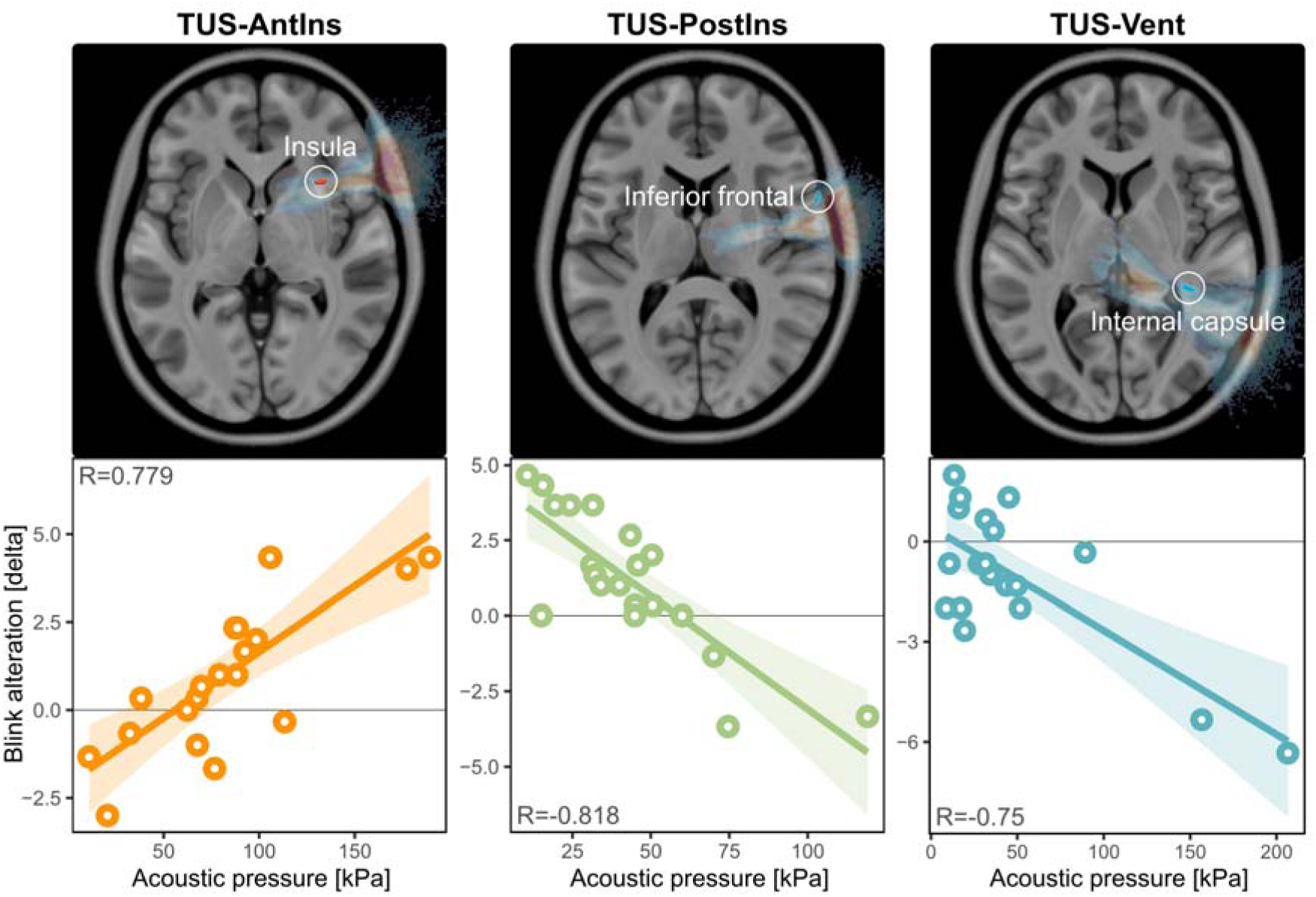
MRI-related effects. For each target, the figure highlights the significant (uncorrected p<0.001) clusters for which the alteration in the number of blinks was found to be significantly associated with the delivered acoustic energy (during TUS for TUS-AntIns and TUS-Vent, 9 minutes following TUS for TUS-PostIns). The bottom part shows the correlations between the blink alteration and the acoustic pressure at the clusters’ peak coordinates.

## Discussion

The present study utilised TUS to explore the causal roles of the anterior and posterior insular cortices in the regulation of action control and urge. First, we confirmed prior findings by demonstrating that the anterior insula plays a role in action control, as evidenced by acute blink suppression alteration when TUS was activated. This effect seems specific since it was not related to a change in the average urge. Second, we observed a dual and delayed role of the posterior insula, with an increase in both the number of blinks and the averaged urge approximately 10 minutes after TUS was applied. Third, and unexpectedly, we discovered an acute effect of TUS-Vent, which decreased the number of blinks. Voxel-based analyses suggest that this effect may have resulted from inadvertently stimulating the PLIC.

### Strengths and limitations

Several aspects should be considered to enhance the interpretability of our findings. First, although the total number of participants (n=20) may appear modest, the use of a within-subject design ensured that all participants completed all experimental conditions with a minimal inter and intra-subject variability. This is supported by the organisation of sessions which were scheduled at the same time of day and, in most cases, on the same day of the week. This aspect strengthened the reliability of our observations.

Second, identical TUS parameters were applied across all participants and targets. While this standardisation ensured methodological consistency, it also introduced variability in the effectively delivered energy due to individual differences in skull density, target depth, and acoustic coupling. Nevertheless, this inter-individual variability provided a valuable opportunity to perform voxel-based analyses linking local acoustic energy to behavioural outcomes, thereby offering additional mechanistic insights.

Third, although a bone-conducting headphone was used to mask the sound produced during TUS, participants were still able to identify the stimulation above chance level. While this represents a limitation, it is important to note that no commonly accepted sham condition currently exists for TUS. Nevertheless, the distinct effects observed across the three targeted regions and the inclusion of the TUS identification score as a covariate in the statistics, strengthen our confidence that the reported effects are due to TUS itself rather than to confounding sensory artefacts. More precise targeting of ultrasound energy is currently only achievable with MRI-guided systems, which unfortunately remain scarce [29].

Finally, data were analysed using robust statistical approaches, including mixed-effects models and FDR corrections for multiple comparisons. These methods increase confidence in the validity of the reported effects but may also have led to the exclusion of potentially meaningful effects that did not reach statistical significance.

### The anterior insula is specifically involved in action control

Applying TUS to the anterior part of the insula induced an acute alteration in action control, reflected by an increased number of blinks. This effect was specific, temporally restricted, and supported by voxel-based analyses showing that higher delivered acoustic energy within the anterior insula was associated with a greater increase in blinks, without any concomitant change in urge. These findings provide several insights: i) they confirm the involvement of the right anterior insula in action control; ii) they indicate a dissociation between action control and the urge-to-blink; iii) they suggest that TUS impaired performance rather than facilitating it; and iv) they indicate that TUS effects on action control are acute.

As expected, our results align with previous neuroimaging evidence implicating the anterior insula in voluntary motor inhibition. Increased activation of the right anterior insula has been reported during blink suppression tasks [7,10], whereas the urge-to-blink has been more closely linked to posterior insula activity [10]. Functionally, the anterior insula is thought to act as a hub integrating interoceptive and exteroceptive information [30] to guide action initiation [7].

The observed increase in blink rate following anterior insula stimulation could be interpreted in several ways. One possibility is that TUS transiently enhanced anterior insula excitability (as supported by TUS studies combined with motor cortex excitability measurement [31]), thereby increasing the likelihood of action initiation. Alternatively, TUS might have reduced the threshold of interoceptive information required to trigger a movement, leading to a higher production of blinks, an approach which could be supported by Bayesian hypothesis [32]. In both cases, TUS appears to have impaired the internal mechanism regulating the urge threshold for blink production.

The exact neurophysiological effects of TUS remain however a matter of debate. While the stimulation parameters used here have been frequently applied in the literature, interpretations of the observed outcomes vary across studies. Some suggest that it corresponds to an excitatory protocol as supported by an increased cortical excitability [31,33]. Others suggest that TUS decreases GABA concentrations, leading to enhanced functional connectivity [18]. Conversely, some suggest that TUS primarily introduces noise rather than a directional modulatory effect [14,34]. Moreover, animal studies have demonstrated that TUS effects depend on tissue composition and cytoarchitecture [35]. This last point is particularly relevant for translation, as we could hypothesise that applying the same parameters to the anterior insula of TS patients, where connectivity patterns are known to be altered [36], could produce different outcomes, keeping the anterior insula and the used TUS parameters a relevant approach to decrease tic frequency.

### The posterior insula has an unclear role in both action control and urge-to-blink

The results obtained for the posterior insula are less straightforward. No acute TUS effect was observed, but both blink frequency and urge ratings increased approximately 10 minutes after the stimulation. Furthermore, the increase in blink number correlated with the delivered acoustic energy within the right IFG and not within the posterior insula. Notably, this correlation was negative, with a higher acoustic pressure associated with fewer blinks.

Although the IFG was not an intended target, its involvement is consistent with its well-established role in inhibitory control and action suppression [37,38], including in TS [39]. The right IFG is a key node in the network subserving response inhibition, particularly in paradigms such as the stop-signal task [40]. Importantly, a recent TUS study targeting the IFG also reported a delayed reduction in functional connectivity with sensorimotor regions [14], suggesting that the delayed behavioural effects observed here could arise from secondary network-level changes rather than direct posterior insula modulation. Together, these findings reinforce the instrumental role of the right IFG in regulating action control.

Interestingly, the delayed increase in blink frequency was accompanied by an increase in the averaged urge, indicating that the coupling between the subjective sensation and the motor act was preserved. This pattern differs from the anterior insula findings, where TUS disrupted the urge-action balance. One interpretation is that TUS applied to the posterior insula primarily modulated interoceptive processing, thereby increasing perceived urge intensity, which in turn elevated blink frequency. This interpretation aligns with previous reports of reduced interoceptive accuracy in individuals with TS compared with healthy controls [41]. Supporting this hypothesis, a recent TUS study comparing stimulation of the anterior versus posterior insula during a pain-rating task demonstrated a selective involvement of the posterior insula in pain intensity perception [42]. These findings suggest that the posterior insula contributes more directly to the integration of interoceptive information, whereas the anterior insula plays a more active role in initiating or inhibiting actions.

### The internal capsule could be a candidate to improve action control

Although stimulation of the posterior ventricle was expected to have no behavioural consequence, we observed an acute reduction in blink frequency when TUS was applied. Post-hoc analyses revealed that this decrease was strongly correlated with the delivered acoustic energy within a cluster located in the PLIC. This tract has been repeatedly reported as abnormal in TS [43–45], with studies showing altered white matter integrity and connectivity within the corticospinal and corticobulbar projections [46,47]. Although the PLIC is not a conventional target for neurosurgical interventions, lesioning or stimulation of adjacent regions such as the PLIC and the ventral striatum has been shown to result in marked tic improvement [48,49].

The PLIC contains major sensorimotor pathways, including corticospinal and corticobulbar fibres that mediate motor output [50,51]. The absence of any concurrent change in urge suggests that TUS may have acutely modulated motor pathway excitability, improving participants’ ability to suppress blinks without altering the subjective difficulty of the task.

Although preliminary, this finding is particularly noteworthy as it represents the only effect observed that points toward an improvement in action control (fewer blinks) and may therefore highlight a potential target for future TUS-based neuromodulation strategies in TS.

## Conclusion

This study provides the first causal evidence that TUS can modulate blink suppression behaviour. Stimulation of the anterior insula impaired action control without altering urge, confirming its critical role in action control. In contrast, posterior insula stimulation produced delayed effects consistent with altered interoceptive processing, while ventricular stimulation unexpectedly improved action control, possibly through modulation of motor fibres within the PLIC. Future studies in TS should determine whether these effects follow the same direction in patients and whether specific stimulation parameters could induce lasting reductions in tic expression.

## Acknowledgements

Stephen Jackson and Aikaterini Gialopsou were funded by MRC Grant (T032588). Stephen Jackson and Aikaterini Gialopsou were funded by a Project grant from Parkinson’s UK (H-2301). Stephen Jackson was funded by an MRC Programme grant (UKRI 527). Stephen Jackson and Mairi Houlgreave were funded by NIHR-funded Nottingham Biomedical Research Centre. Cyril Atkinson-Clement and Caitlin Smith were supported by a Postdoctoral Research Fellowship funded by a philanthropic donation from Daniel Katz Ltd.

## Author contributions

**Cyril Atkinson-Clement**: Conceptualisation, Methodology, Formal analysis, Investigation, Data Curation, Writing - Original Draft, Visualisation. **Mairi Houlgreave**: Conceptualisation, Methodology, Writing - Review & Editing. **Aikaterini Gialopsou**: Conceptualisation, Methodology, Writing - Review & Editing. **Caitlin Mairi Smith**: Writing - Review & Editing.

**Isabel Farr**: Writing - Review & Editing. **Aneta Dvorakova**: Writing - Review & Editing. **James Kennaway**: Writing - Review & Editing. **Stephen R. Jackson**: Conceptualisation, Methodology, Writing - Review & Editing, Supervision, Project administration, Funding acquisition.

## Data availability statement

All data and original code used to analyse the data in this paper are available from the corresponding contact upon request.

**Figure.sup1.** Anterior and posterior parts of the insula. The figure shows the insula location based on the AAL atlas [25], with the anterior part highlighted in blue and the posterior part highlighted in red. The spheres correspond to the centre of each of these regions of interest and were targeted with TUS.

**Figure.sup2.** Illustration of the different steps obtained for one participant. From top to bottom, it corresponds to identify the target location (obtain by co-registering the native T1 with the MNI template), to determine the transducer position which allow to get the lower head-transducer angle [27], to estimate a PCT derived from the T1 [28], to segment the tissues to determine the sound speed (achieved by k-Plan), to determine the delivered acoustic energy and the temperature increase based on the used TUS parameters (achieved by k-Plan).

## References

1. Diagnostic and Statistical Manual of Mental Disorders: DSM-5. (American Psychiatric Publishing, Washington, DC, 2013).

2. Sambrani, T., Jakubovski, E. & Müller-Vahl, K. R. New Insights into Clinical Characteristics of Gilles de la Tourette Syndrome: Findings in 1032 Patients from a Single German Center. Front Neurosci 10, 415 (2016).

3. Robertson, M. M. Gilles de la Tourette syndrome: the complexities of phenotype and treatment. Br J Hosp Med 72, 100–S7 (2011).

4. Wilhelm, S. et al. Randomized Trial of Behavior Therapy for Adults With Tourette Syndrome. Arch Gen Psychiatry 69, 795 (2012).

5. Bohlhalter, S. Neural correlates of tic generation in Tourette syndrome: an event-related functional MRI study. Brain 129, 2029–2037 (2006).

6. Jackson, S. R., Parkinson, A., Kim, S. Y., Schüermann, M. & Eickhoff, S. B. On the functional anatomy of the urge-for-action. Cognitive Neuroscience 2, 227–243 (2011).

7. Berman, B. D., Horovitz, S. G., Morel, B. & Hallett, M. Neural correlates of blink suppression and the buildup of a natural bodily urge. NeuroImage 59, 1441–1450 (2012).

8. Botteron, H. E. et al. The urge to blink in Tourette syndrome. Cortex 120, 556–566 (2019).

9. Zouki, J.-J. et al. Functional brain networks associated with the urge for action: Implications for pathological urge. Neuroscience & Biobehavioral Reviews 163, 105779 (2024).

10. Houlgreave, M. S. et al. Uncovering the neural correlates of the urge-to-blink: A study utilising subjective urge ratings and paradigm free mapping. Imaging Neuroscience 3, IMAG.a.84 (2025).

11. Jackson, S. R. et al. The role of the insula in the generation of motor tics and the experience of the premonitory urge-to-tic in Tourette syndrome. Cortex 126, 119–133 (2020).

12. Sarica, C. et al. Human Studies of Transcranial Ultrasound neuromodulation: A systematic review of effectiveness and safety. Brain Stimulation 15, 737–746 (2022).

13. Atkinson-Clement, C. et al. Dynamical and individualised approach of transcranial ultrasound neuromodulation effects in non-human primates. Sci Rep 14, 11916 (2024).

14. Atkinson-Clement, C., Alkhawashki, M., Gatica, M., Ross, J. & Kaiser, M. Dynamic changes in human brain connectivity following ultrasound neuromodulation. Sci Rep 14, 30025 (2024).

15. Atkinson-Clement, C. et al. Temporal dynamics of offline transcranial ultrasound stimulation. Current Research in Neurobiology 8, 100148 (2025).

16. Darmani, G. et al. Non-invasive transcranial ultrasound stimulation for neuromodulation. Clinical Neurophysiology 135, 51–73 (2022).

17. Bault, N., Yaakub, S. N. & Fouragnan, E. Early-Phase Neuroplasticity Induced by Offline Transcranial Ultrasound Stimulation in Primates. https://osf.io/grakn (2023) doi:10.31234/osf.io/grakn.

18. Yaakub, S. N. et al. Transcranial focused ultrasound-mediated neurochemical and functional connectivity changes in deep cortical regions in humans. Nat Commun 14, 5318 (2023).

19. Yoo, S., Mittelstein, D. R., Hurt, R. C., Lacroix, J. & Shapiro, M. G. Focused ultrasound excites cortical neurons via mechanosensitive calcium accumulation and ion channel amplification. Nat Commun 13, 493 (2022).

20. Plaksin, M., Kimmel, E. & Shoham, S. Cell-Type-Selective Effects of Intramembrane Cavitation as a Unifying Theoretical Framework for Ultrasonic Neuromodulation. eneuro **3**, ENEURO.0136-15.2016 (2016).

21. Oh, S.-J. et al. Ultrasonic Neuromodulation via Astrocytic TRPA1. Current Biology 29, 3386–3401.e8 (2019).

22. Lee, K., Park, T. Y., Lee, W. & Kim, H. A review of functional neuromodulation in humans using low-intensity transcranial focused ultrasound. Biomed. Eng. Lett. 14, 407– 438 (2024).

23. World Medical Association General Assembly. Declaration of Helsinki, Amendment. (2004).

24. Kop, B. R., et al. Auditory confounds can drive online effects of transcranial ultrasonic stimulation in humans. Preprint at 10.7554/eLife.88762.2 (2024).

25. Tzourio-Mazoyer, N. et al. Automated anatomical labeling of activations in SPM using a macroscopic anatomical parcellation of the MNI MRI single-subject brain. Neuroimage 15, 273–289 (2002).

26. Avants, B., Tustison, N. J. & Song, G. Advanced Normalization Tools: V1.0. The Insight Journal 10.54294/uvnhin (2009) doi:10.54294/uvnhin.

27. Atkinson-Clement, C. & Kaiser, M. Optimizing Transcranial Focused Ultrasound Stimulation: An Open-source Tool for Precise Targeting. Neuromodulation: Technology at the Neural Interface S1094715924006275 (2024) doi:10.1016/j.neurom.2024.06.496.

28. Yaakub, S. N. et al. Pseudo-CTs from T1-weighted MRI for planning of low-intensity transcranial focused ultrasound neuromodulation: An open-source tool. Brain Stimulation 16, 75–78 (2023).

29. Martin, E. et al. Ultrasound system for precise neuromodulation of human deep brain circuits. Nat Commun 16, 8024 (2025).

30. Chang, L. J., Yarkoni, T., Khaw, M. W. & Sanfey, A. G. Decoding the Role of the Insula in Human Cognition: Functional Parcellation and Large-Scale Reverse Inference. Cerebral Cortex 23, 739–749 (2013).

31. Zeng, K. et al. Effects of different sonication parameters of theta burst transcranial ultrasound stimulation on human motor cortex. Brain Stimulation 17, 258–268 (2024).

32. Rae, C. L., Critchley, H. D. & Seth, A. K. A Bayesian Account of the Sensory-Motor Interactions Underlying Symptoms of Tourette Syndrome. Front. Psychiatry 10, 29 (2019).

33. Xia, X. et al. Effects of the motor cortical theta burst transcranial[focused ultrasound stimulation on the contralateral motor cortex. The Journal of Physiology 602, 2931–2943 (2024).

34. Gatica, M. et al. Understanding the high-order network plasticity mechanisms of ultrasound neuromodulation. PLoS Comput Biol 21, e1013514 (2025).

35. Tseng, H. et al. Region-specific effects of ultrasound on individual neurons in the awake mammalian brain. iScience 24, 102955 (2021).

36. Tinaz, S., Malone, P., Hallett, M. & Horovitz, S. G. Role of the right dorsal anterior insula in the urge to tic in tourette syndrome: Right Dorsal Anterior Insula in Tourette. Mov Disord. 30, 1190–1197 (2015).

37. Aron, A. R. Cortical and Subcortical Contributions to Stop Signal Response Inhibition: Role of the Subthalamic Nucleus. Journal of Neuroscience 26, 2424–2433 (2006).

38. Aron, A. R., Robbins, T. W. & Poldrack, R. A. Inhibition and the right inferior frontal cortex: one decade on. Trends in Cognitive Sciences 18, 177–185 (2014).

39. Atkinson-Clement, C. et al. Neural correlates and role of medication in reactive motor impulsivity in Tourette disorder. Cortex 125, 60–72 (2020).

40. Aron, A. R. & Poldrack, R. A. The Cognitive Neuroscience of Response Inhibition: Relevance for Genetic Research in Attention-Deficit/Hyperactivity Disorder. Biological Psychiatry 57, 1285–1292 (2005).

41. Rae, C. L., Larsson, D. E. O., Garfinkel, S. N. & Critchley, H. D. Dimensions of interoception predict premonitory urges and tic severity in Tourette syndrome. Psychiatry Research 271, 469–475 (2019).

42. In, A., Strohman, A., Payne, B. & Legon, W. Low-intensity focused ultrasound to the posterior insula reduces temporal summation of pain. Brain Stimulation 17, 911–924 (2024).

43. Wen, H. et al. Combining tract[and atlas[based analysis reveals microstructural abnormalities in early Tourette syndrome children. Human Brain Mapping 37, 1903–1919 (2016).

44. Neuner, I. et al. White-matter abnormalities in Tourette syndrome extend beyond motor pathways. NeuroImage 51, 1184–1193 (2010).

45. Bruce, A. B. et al. Altered frontal-mediated inhibition and white matter connectivity in pediatric chronic tic disorders. Exp Brain Res 239, 955–965 (2021).

46. Worbe, Y. et al. Altered structural connectivity of cortico-striato-pallido-thalamic networks in Gilles de la Tourette syndrome. Brain 138, 472–482 (2015).

47. Greene, D. J., Schlaggar, B. L. & Black, K. J. Neuroimaging in Tourette Syndrome: Research Highlights from 2014 to 2015. Current Developmental Disorders Reports 2, 300–308 (2015).

48. Sun, B., Krahl, S. E., Zhan, S. & Shen, J. Improved Capsulotomy for Refractory Tourette’s Syndrome. Stereotact Funct Neurosurg 83, 55–56 (2005).

49. Baldermann, J. C. et al. Deep Brain Stimulation for Tourette-Syndrome: A Systematic Review and Meta-Analysis. Brain Stimulation 9, 296–304 (2016).

50. Qian, C. & Tan, F. Internal capsule: The homunculus distribution in the posterior limb. Brain and Behavior 7, e00629 (2017).

51. Forhad, C., Mohammod, H., Mainul, S., Shamim, A. & Mohammod, I. White fiber dissection of brain; the internal capsule: a cadaveric study. Turkish Neurosurgery 10.5137/1019-5149.JTN.3052-10.2 (2010) doi:10.5137/1019-5149.JTN.3052-10.2.

