## Supplementary figures and images for "Disrupting action control with transcranial ultrasound neuromodulation: a step forward for Tourette syndrome"

### Figure.sup1

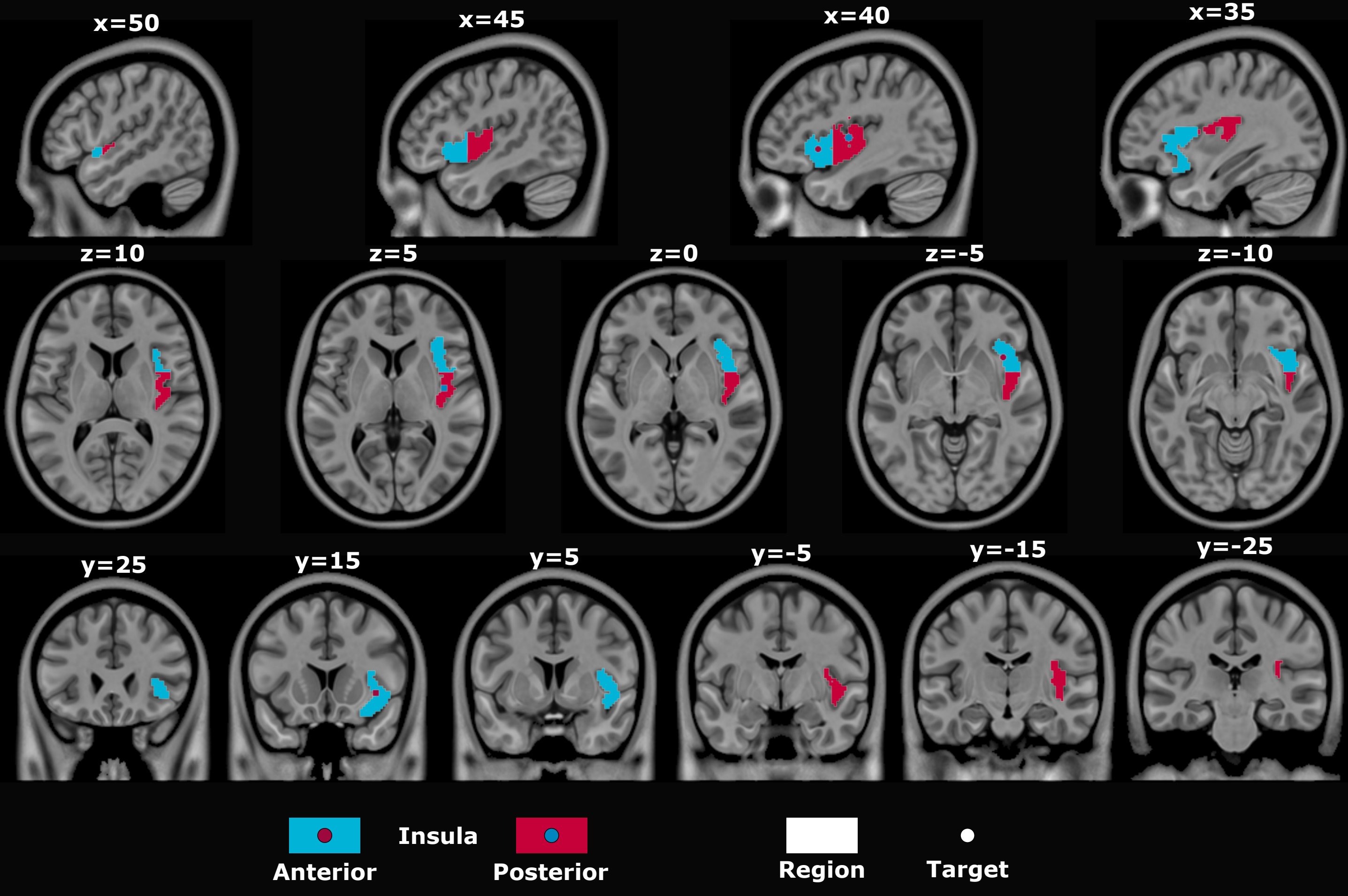

### Figure.sup2

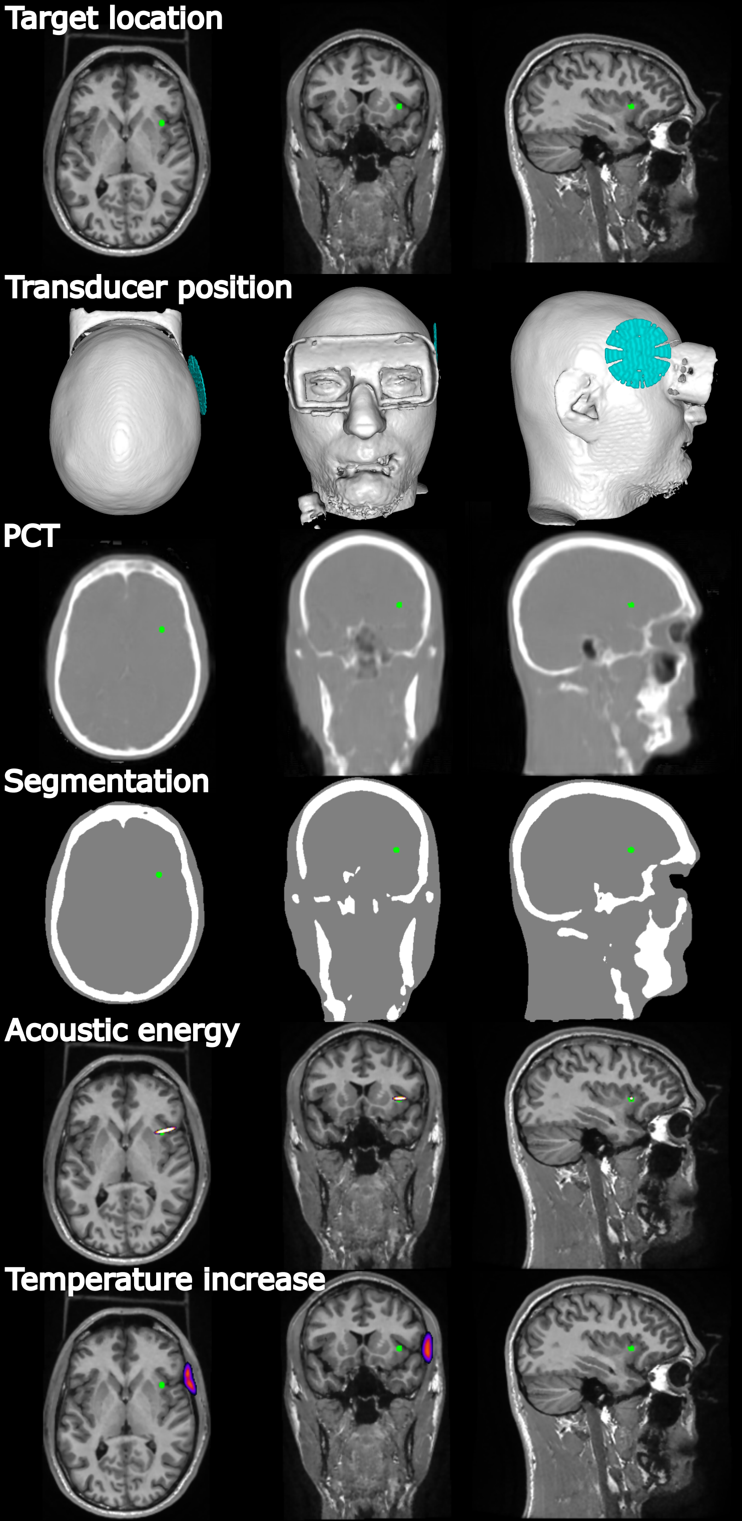
